# Effects of gypenosides on enteroendocrine L-cell function and GLP-1 secretion

**DOI:** 10.1101/2020.09.07.285916

**Authors:** Chinmai Patibandla, Erin Campbell, Xinhua Shu, Angus M Shaw, Sharron Dolan, Steven Patterson

## Abstract

Glucagon-like peptide 1 (GLP-1) is an incretin hormone produced in gut L-cells, which regulates postprandial glucose-dependent insulin secretion, also known as the incretin effect. GLP-1 secretion may be reduced in type 2 diabetes mellitus, impacting on glycaemic regulation. Thus, methods to enhance endogenous GLP-1 secretion by use of natural GLP-1 secretagogues may improve glucose control in diabetes. Gypenosides (GYP) extracted from the plant *Gynostemma Pentaphyllum* (Jiaogulan) are known for their glucose-lowering effects both *in vitro* and *in vivo*, although their effect on GLP-1 secretion is unknown. Our results showed that GYP enhanced cell viability and significantly upregulated antioxidant gene Nrf2, Cat and Ho-1 expression. GYP did not affect glucokinase expression but downregulated proglucagon gene expression over 24h, although, cellular GLP-1 content was unchanged. Prohormone convertase 1 (Pcsk1) gene expression was unchanged by GYP over 24h, although protein levels were significantly downregulated, while prohormone convertase 2 (Pcsk2) mRNA and protein levels were significantly upregulated. Acute exposure to gypenosides enhanced calcium uptake and GLP-1 release from GLUTag cells both at low and high glucose concentrations. These results suggest that anti-diabetic properties of gypenosides are partly linked to their ability to stimulate GLP-1 secretion. Gypenosides enhance antioxidant gene expression and may protect L-cells from excess oxidative stress.

## 1 Introduction

Glucagon-like peptide-1 (GLP-1) is an intestinal hormone secreted by enteroendocrine L-cells. The proglucagon gene is expressed in L-cells as well as pancreatic alpha cells, and post-translational cleavage by prohormone convertase-1 (Pcsk1) gives rise to peptides glicentin, oxyntomodulin, GLP-1 and GLP-2 (Dhanvantari et al., 1996; Rouillé et al., 1995). Secreted GLP-1 (1-37) exists in two bioactive forms, GLP-1 (7-36), GLP-1 (7-37) amide but is rapidly degraded by dipeptidyl peptidase-IV (DPP-IV) to GLP-1 (9-36) and GLP-1 (9-37) amide (Hansen et al., 1999). Active GLP-1 has many systemic functions including, glucose homeostasis by promoting glucose-dependent postprandial insulin secretion from pancreatic β-cells, enhancing insulin gene expression, β-cell proliferation, activating protective anti-apoptotic pathways and inhibiting the release of glucagon from islet α-cells (Drucker, 2006). It has been reported that GLP-1 secretion is reduced along with the impairment of the incretin effect in type 2 diabetes mellitus (T2DM) (Mannucci et al., 2000).

Hyperglycaemic- and hyperlipidaemic-induced pancreatic β-cell oxidative stress and apoptosis in T2DM has been well studied *in vivo* and *in vitro* but similar studies in L-cells are scarce due to their location and distribution in the gut. Previous studies in L-cell model (GLUTag cells) showed that chronic exposure to palmitate and glucose, significantly reduced cell viability (Vasu *et al.*, 2015), and altered proglucagon gene processing (Hayashi et al., 2014), indicating that hyperglycemia and hyperlipidaemia might also have a detrimental effect on L-cells and perhaps contribute to impaired incretin responses. Activation of the Nrf2 antioxidant pathway has significant protective effects against oxidative stress in pancreatic β-cells (Abebe et al., 2017; Masuda et al., 2015). A similar Nrf2 mediated protective effect may also protect L-cells exposed to prolonged stress. Thus, any pharmacological agents stimulating endogenous GLP-1 secretion from gut L-cells that also possess antioxidant mediated cell protection could prove beneficial in the management of T2DM.

*Gynostemma pentaphyllum* (GP) Makino, also known as Jiaogulan, grows in several parts of Asia (China, Japan, Vietnam, India and Korea) and has been used in Chinese folk medicine for centuries due to its proposed wide-ranging health benefits. Anti-diabetic properties of GYP are well documented both in T2DM patients and diabetic rodent models. While the majority of previous studies on the anti-diabetic properties of GYP have focused on β-cells and insulin secretion, hepatic glucose output (Yassin et al., 2011) and adipose insulin sensitivity (Liu et al., 2017), its effect on gut endocrine function, specifically GLP-1 secreting L-cells has yet to be investigated. Thus the current study is focused on investigating GYP effects on GLP-1 secretion and L-cell function using the robustly characterised mouse intestinal L-cell model, GLUTag cell line (Drucker et al., 1994; Kuhre et al., 2016).

## 2 Methods

### 2.1 Cell Culture and viability

GLUTag cells (a kind gift from Prof. D Drucker) were routinely cultured in Dulbecco’s modified eagle medium (DMEM) (5.5 mmol/L D-glucose) (Lonza, UK), supplemented with 10% (v/v) foetal bovine serum (v/v), 50U/ml penicillin/streptomycin and 2mM L-glutamine, and were maintained at 37°C with 5% CO_2_ and 95% air (Drucker et al., 1992; Lee et al., 1992). Cells were trypsinised and sub-cultured at 1:5 dilutions when 80-90% confluence was reached. Passage numbers from 25 to 35 were used for experiments. To determine the effects of CPE on cell viability over 24h, GLUTag cells were seeded in Geltrex™ (Gibco, UK) coated 96 well plates at 10,000 cells/well. After overnight culture, cells were incubated with CPE (0.78 to 200μg/ml) for 24h and cell viability assessed using MTT assay.

MIN6 cells were cultured in DMEM containing 4.5g/L glucose supplemented with 10% FBS (v/v) and 50U/ml penicillin/streptomycin. Cells were maintained at 37°C with 5% CO_2_ and 95% air. When 80-90% confluence was reached, cells were detached from the flask using trypsin/EDTA and sub-cultured at 1:3 dilution.

### 2.2 GYP Extract

Gypenosides were purchased from Xi’an Jiatian Biotech Co. Ltd, China (purity 98%). Gypenosides were dissolved in absolute ethanol (25mg/ml) by continuous shaking at room temperature overnight and stored at −20°C until use.

### 2.3 Ca^2+^ measurement

GLUTag cells were seeded on Geltrex™ coated glass coverslips and cultured until confluent. Prior to use, cells were incubated for 45 min with 2μM FURA-2AM (Tocris, UK) in KRB Buffer containing 1.1 mM glucose at 37°C. The KRB buffer was composed of (in mmol/L): 115 NaCl, 4.7 KCl, 1.2 KH_2_PO_4_, 1.2 MgSO_4_, 1.28 CaCl_2_, 20 HEPES and 0.1% (w/v) BSA (pH 7.4). Coverslips were rinsed with PBS and mounted onto an RC-21BRW closed bath imaging chamber (Warner Instruments) with P-2 platform and cells were perfused at a rate of 1ml/min with KRB Buffer with/without GYP. A Nikon Eclipse TE2000-U microscope fitted with Photometrics Cool SNAP™ HQ Camera was used to acquire images. Fluorescence emission was recorded and cytosolic Ca^2+^ changes were plotted as 340nm/380nm ratio over time.

### 2.4 Measurement of GLP-1 Secretion by ELISA

For secretion tests, cells were seeded on 12 well plates (2.5 × 10^5^ cells/well) coated with Geltrex™ and incubated for 24h. Prior to use, cells were washed and incubated for 30 mins with KRB Buffer (1.1mM Glucose) followed by 1h incubation with test reagents. The supernatant was collected and centrifuged at 900g for 5mins and GLP-1 secretion over 1h treatment with GYP was measured using a total GLP-1 ELISA kit (Millipore, UK) according to manufacturer’s specifications.

### 2.5 Immunofluorescence

To analyse GLP-1 staining, GLUTag cells (1X10^5^ cells/ coverslip) were seeded on Geltrex™ coated glass coverslips and cultured overnight until confluent. After treatment with 100μg/ml GYP for 24h, cells were fixed with 4% paraformaldehyde for 10 mins at 4°C followed by 30mins blocking with 2% BSA and stained with anti-GLP-1 antibody (1:200) (Gt pAb to GLP-1, SC-26637, Santa Cruz Biotechnology) overnight in a humid chamber at 4°C. The secondary antibody was added and incubated at room temperature for 1h (Alexafluor 594 donkey anti-goat IgG secondary antibody (1:500) Abcam, ab150132). Mounting medium with DAPI (Abcam, ab104139) was used to stain the nucleus and slides were visualised and images were captured using EVOS FL imaging system.

### 2.6 RNA extraction and PCR

Total RNA was extracted from GYP treated GLUTag cells using the NucleoSpin® RNA kit (Macherey-Nagel, UK) according to the manufacturer’s protocol. From total RNA, cDNA was synthesised using High-Capacity cDNA reverse transcription kit (Applied Biosystems, UK). Using mouse Ncx1, Ncx2 and Ncx3 specific primers, cDNA was amplified by PCR using 5x FIREPol® Master Mix (Solis-BioDyne, Estonia) and β-Actin was used as the housekeeping gene. Cycling conditions were; initial denaturation at 95°C for 4 mins followed by 40 cycles of 95°C for 20 sec, 54°C for 45 sec, 72°C for 45 sec; and final elongation was at 72°C for 5 mins. Primers used are Ncx1,F-CCTTGTGCATCTTAGCAATG, R-TCTCACTCATCTCCACCAGA; Ncx2, F-CACGCACCTTCCCTGATTTA, R-CTCATCCACTAAGCTGGTTCTC; Ncx3, F-CTATCCCACCACCCAACTATTC, R-GTCAGCCTGGAACACTCTTATC; which yielded product sizes of 437bp, 316bp and 482 bp, respectively. PCR products were separated by 2% agarose gels and visualised by Midori green (Geneflow, UK).

### 2.7 Quantitative real-time PCR

Quantitative real-time PCR was performed on a CFX96™ Real-Time PCR detection system using iQ™ SYBR® Green Supermix (BIO-RAD, UK) according to manufacturer’s specifications. Primer sequences used were listed in table 1.

**Table 1:**
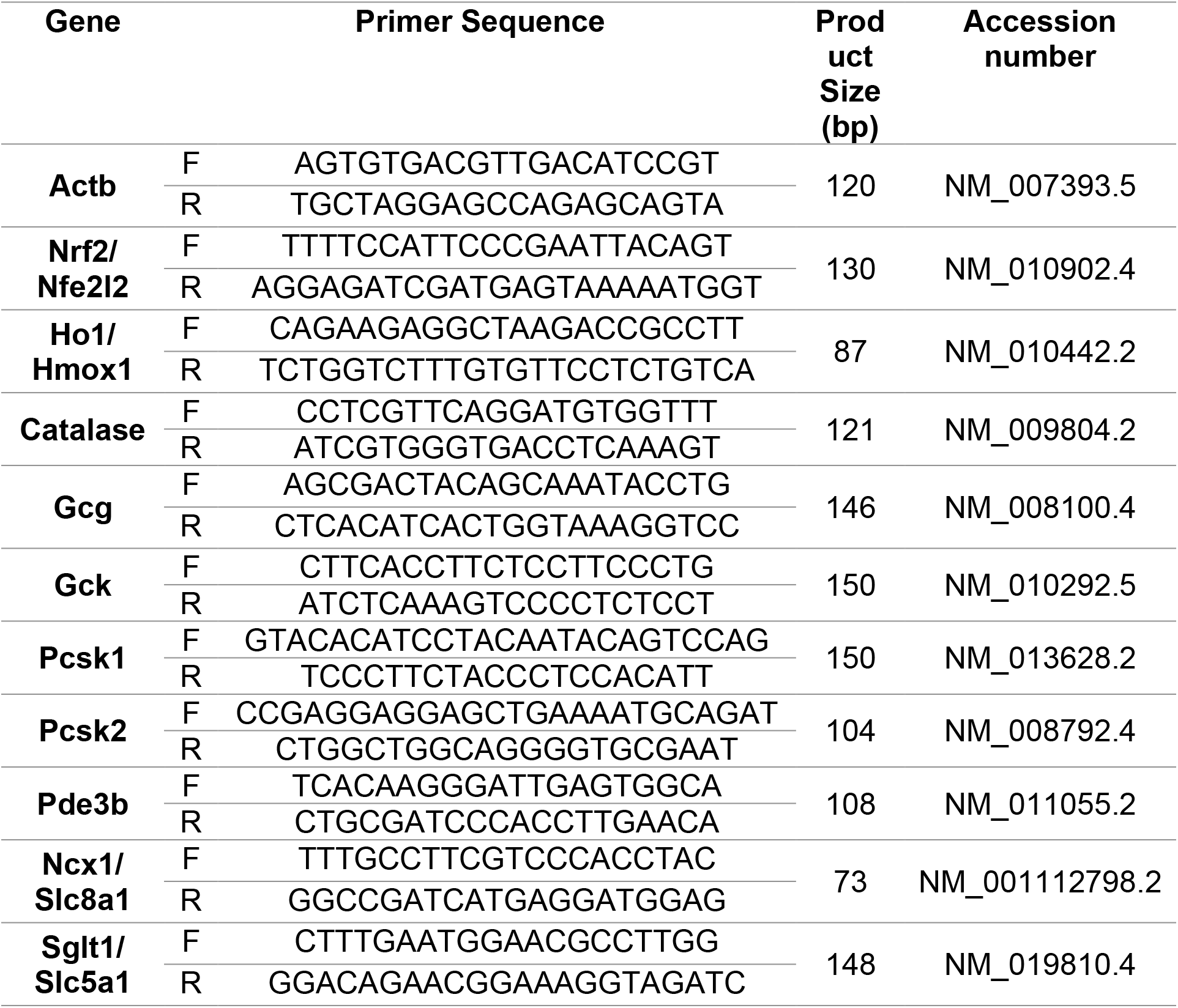
List of mouse primers used in the quantitative real-time PCR

### 2.8 Western blot

Cells were lysed with ice-cold radioimmunoassay precipitation buffer (RIPA) (50mM Tris-HCl pH 8.0, 150mM NaCl, 1% NP-40, 0.5% sodium deoxycholate, 0.1% sodium dodecyl sulphate) and centrifuged at 10,000 rpm for 15mins (4°C) to separate any cell debris. Protein concentration in the supernatant was measured using DC™ Protein assay kit (BIO-RAD, UK) according to manufacturer’s specifications using bovine serum albumin (BSA) as standard. Proteins were separated by SDS-PAGE using hand cast gradient gels (4-12%) and transferred to nitrocellulose membrane (20V for 10mins) using the iBlot® blotting system (Thermo Scientific, UK). Membranes were blocked with 5% (w/v) BSA for 1h at room temperature and incubated with primary antibodies overnight at 4°C. Primary antibodies used were: Anti-Pcsk1 (1:1000) (GTX113797S, Genetex), Anti-Pcsk2 (2μg/ml) (MAB6018-SP, Novus Biologicals) and Anti-Actin-beta (1:1000) (ab-1801, Abcam UK). Blots were incubated with IRDye® conjugated specific secondary antibodies (1:5000) for 1h at room temperature. Signals were detected using the Odyssey® Fc imaging system (LI-COR, UK) and analysed using Image Studio™ software.

### 2.9 Data Analysis

Results were presented as means ± S.E.M. Data were analysed in Graphpad PRISM® software (ver 6.01) by using unpaired student’s t-test (parametric, two-tailed) for comparing two groups or one-way ANOVA for comparing more than 2 groups with Dunnett’s post hoc test with a significance threshold of p<0.05.

## 3 Results

### 3.1 Effects of GYP on GLUTag cell viability

To elucidate the effect of GYP extract on GLUTag cell viability, cells were incubated for 24h with different concentrations (3.125μg/ml – 200μg/ml) of GYP and an MTT assay performed to assess the cell viability. At high concentration of 200μg/ml GYP significantly reduced cell viability (p<0.0001) by 36.4% (Fig.1). Lower concentration of 100, 50 and 25μg/ml enhanced GLUTag viability compared to control (for 100 & 50 μg/ml P<0.0001; for 25 μg/ml P<0.05) (fig.1).

**Figure 1:**
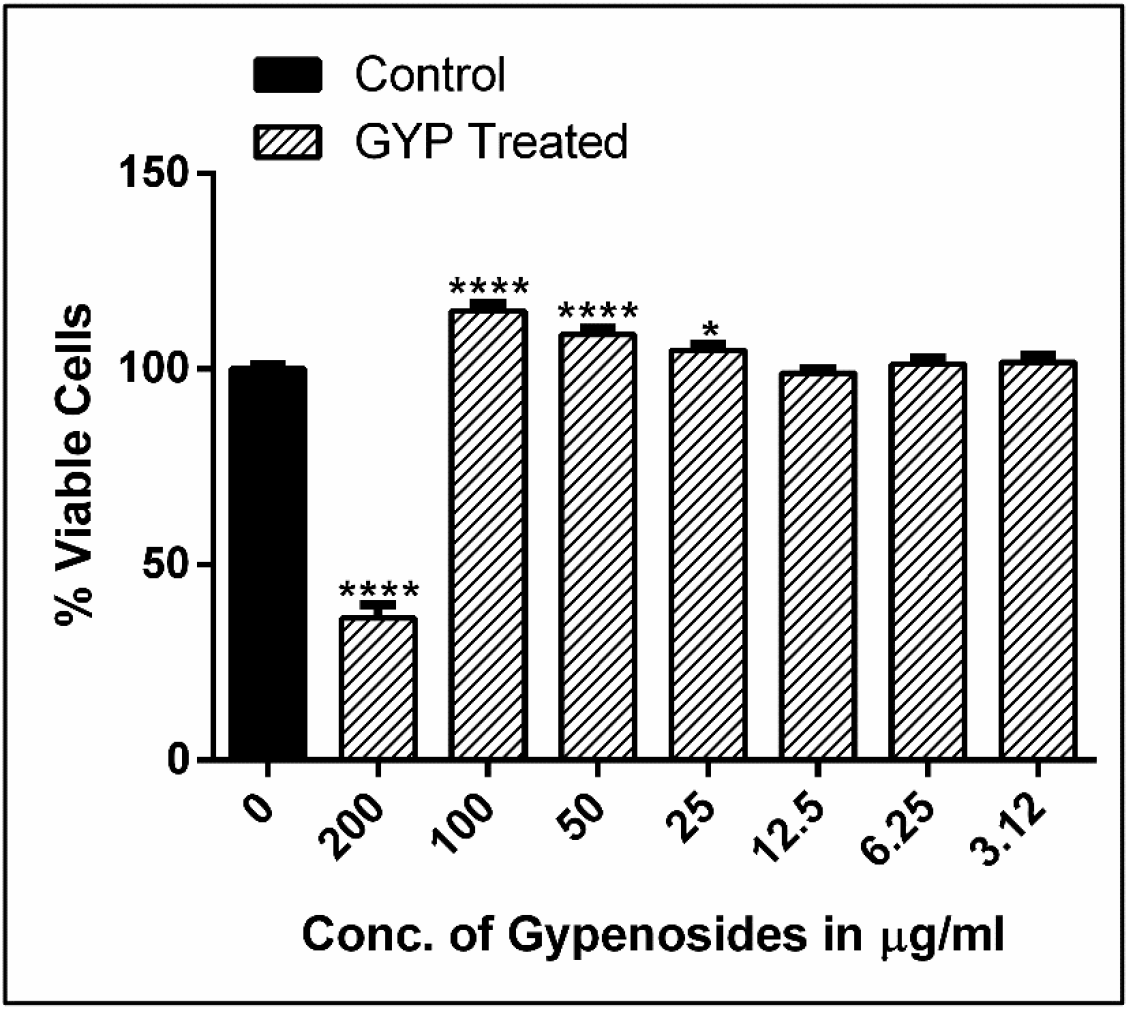
Effects of GYP at concentrations between 3.12 - 200μg/ml on GLUTag cell viability over 24 h treatment. Plotted as % change in cell viability compared to control. Values represent mean ±S.E.M. from three different experiments (n=8). Student’s t test was used for statistical analysis. *, P<0.05; ****, P<0.0001.

### 3.2 Effects of GYP on GLP-1 release and intracellular calcium [Ca^2+^]_i_

GYP treatment over 1h concentration-dependently stimulated GLP-1 secretion from GLUTag cells both at low (1.1mM) and high (16.7mM) glucose concentrations (Fig 2A). At low glucose concentration, 50 & 100 μg/ml GYP increased GLP-1 release 2.7-(P<0.05) and 7.5-fold (P<0.0001), respectively. At high glucose, these concentrations elicited 3.6-(P<0.001) and 13.5-fold (P<0.01) increases in GLP-1 release compared to high glucose alone. There was no significant difference in GLP-1 secretion between low compared to high glucose for any GYP concentration.

**Figure 2:**
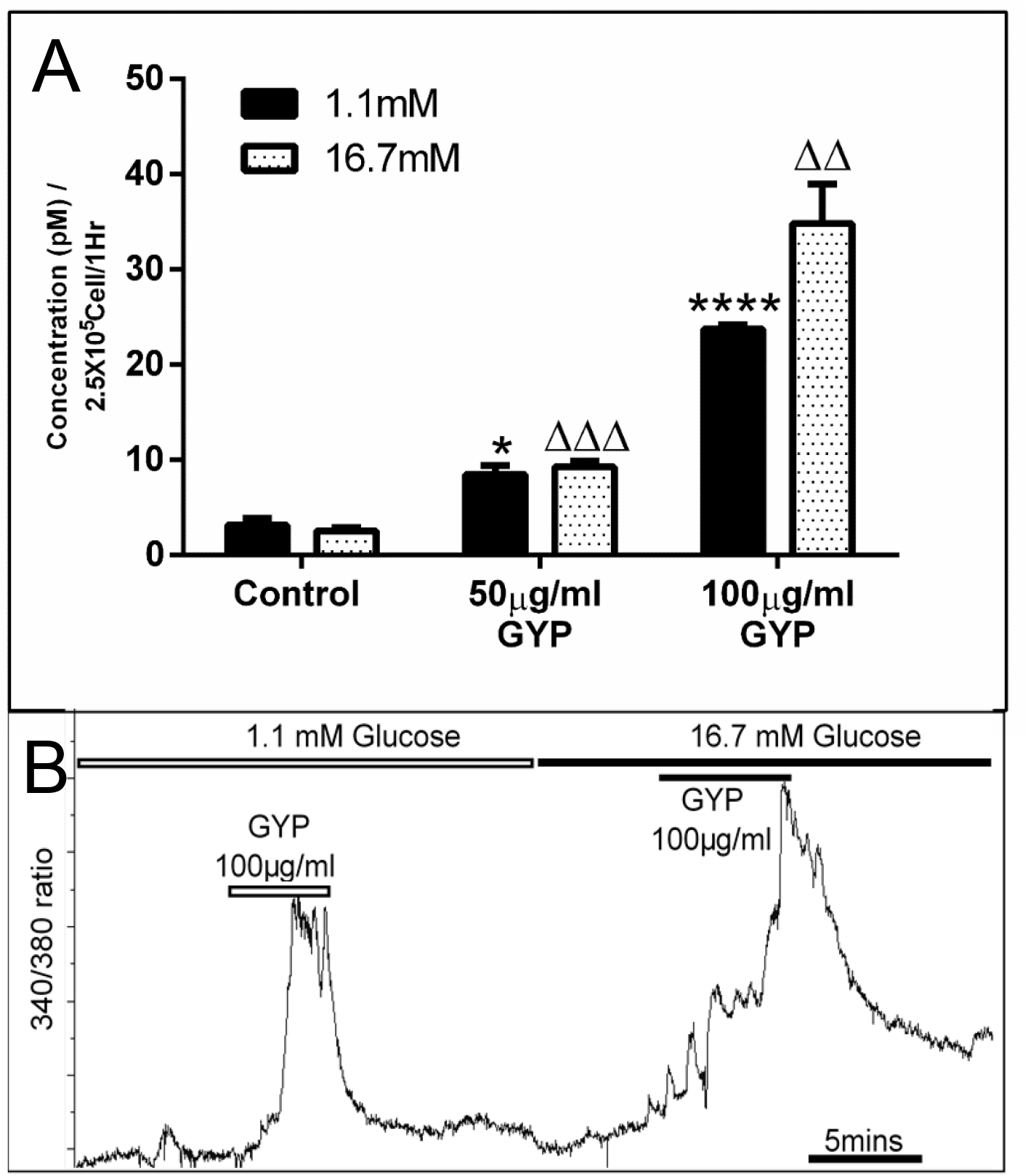
Effects of GYP on GLP-1 secretion and intracellular calcium in GLUTag L-cells. **(A)** GLP-1 secretion measured in presence of GYP 50 & 100μg/ml at low (1.1mM) (Black) or high (16.7mM) (dotted) glucose concentration. *, indicates significance compared to 1.1mM glucose control and Δ, was significance compared to 16.7 glucose control. Values plotted as mean ±S.E.M. from 4 independent experiments conducted in duplicate. *, P<0.05; ****, P<0.0001; ΔΔ, P<0.01 and ΔΔΔ, P<0.001. **(B)** Changes in intracellular calcium levels in GLUTag cells in presence of 100μg/ml GYP at low (1.1mM) and high (16.7) glucose concentration. Response was representative of 44 cells from 3 independent experiments.

As for all endocrine secretory cells, intracellular calcium levels play a fundamental role in GLP-1 exocytosis. GLUTag cells were perfused with GYP in the presence of low and high glucose concentrations. In response to acute exposure to GYP (100μg/ml), [Ca^2+^]^i^ was increased at both low (1.1mM) and high (16.7mM) glucose concentrations, as shown in Figure 2B.

### 3.3 Effects of GYP treatment on L-cell gene expression

Pro-inflammatory transcription factor, nuclear factor -kappa-B p105 (Nfkb1) was unchanged after 24h GYP treatment, whereas Nuclear Factor, Erythroid 2 Like 2 (Nfe2l2 / Nrf2), heme oxygenase 1 (Hmox1/ Ho1) (P<0.001) and catalase (Cat) expression were all increased by GYP (P<0.001 - 0.0001) (Fig. 4.5A). GYP treatment (100μg/ml) over 24h significantly downregulated proglucagon (Gcg) (P<0.001) gene expression in GLUTag cells, although glucokinase (Gck) and prohormone convertase 1 (Pcsk1) expression were unchanged (Fig. 4.5B). Interestingly, prohormone convertase 2 expression was enhanced (P<0.0001) following 24h GYP treatment. Expression of the main regulator of glucose uptake in L-cells, the sodium/glucose cotransporter-1 (Sglt-1/ Slc5a1), was also upregulated (P<0.0001) along with sodium/calcium exchanger Ncx1 (P<0.05) (Fig. 4.5C).

**Figure 3:**
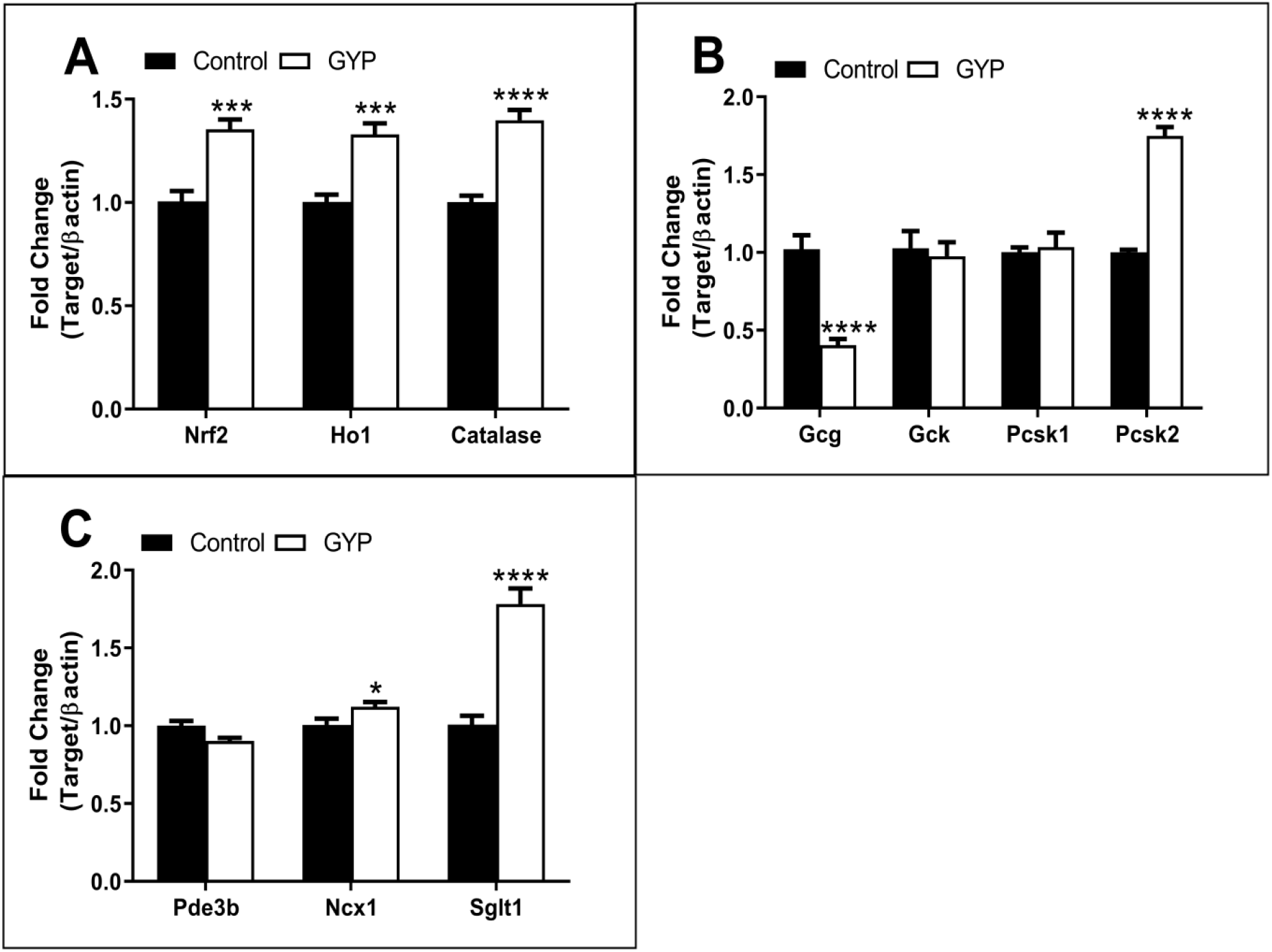
Effects of 24h culture with GYP (100μg/ml) on expression of key genes in GLUTag cell. Data represent fold change in mRNA levels compared to control/untreated GLUTag cell and normalised to β-actin expression. Values represent mean ± S.E.M. from three independent experiments performed in duplicate. Student’s t test was used for statistical analysis. *, P<0.05; ***, P<0.001; ****, P<0.0001.

**Figure 4:**
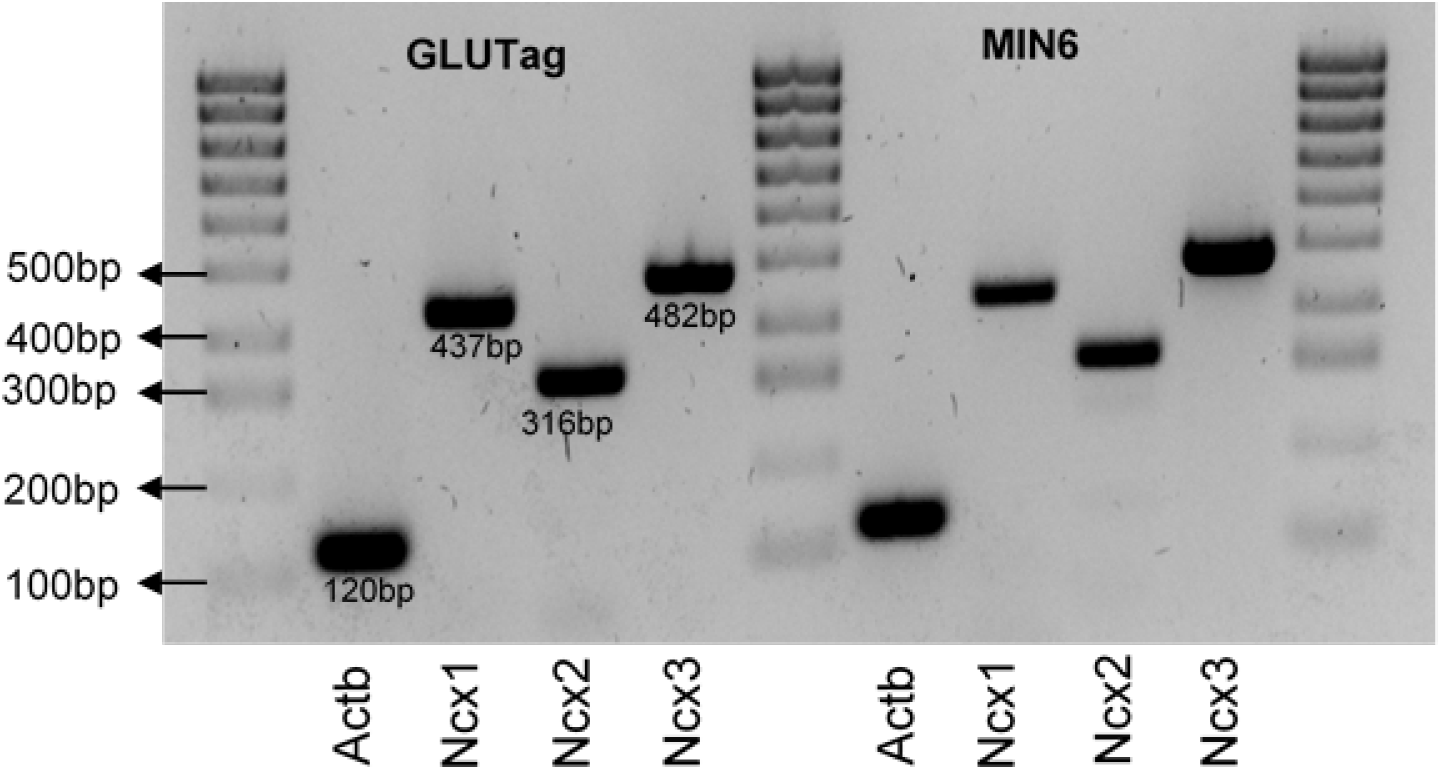
Expression of sodium/calcium exchanger in GLUTag cells and mouse insulinoma MIN6 cells. Amplification of GLUTag mRNA by PCR detected Ncx1, Ncx2 and Ncx3. MIN6 cell mRNA was used as positive control for the primer pairs which are shown on the right.

### 3.4 Expression of Ncx isoforms in GLUTag cells

Although GYP induced a significant increase in Ncx 1 expression, it has not previously been confirmed which other Ncx isoforms are expressed in GLUTag cells. RT-PCR was performed followed by gel electrophoresis to identify the Ncx isoforms. PCR product identity was confirmed by sequencing, confirming the expression of all three isoforms of Ncx (Ncx1, Ncx2 and Ncx3) in GLUTag L-cells (Fig.4). We have used mouse β-cell line MIN6 cell cDNA as a positive control as it is previously known that Ncx isoforms are expressed in pancreatic β-cells (Fig. 4).

### 3.5 Effects of GYP on Protein expression and GLP-1 staining

At a concentration of 100 μg/ml, GYP treatment over 24h enhanced Pcsk2 (P<0.05) and downregulated Pcsk1 (P<0.01) protein levels (Fig.5). As 24h treatment with GYP downregulated proglucagon expression and Pcsk1 protein levels, we analysed the possible changes in staining intensity of cellular GLP-1 by immunostaining and the results showed no significant difference (Fig. 6).

**Figure 5:**
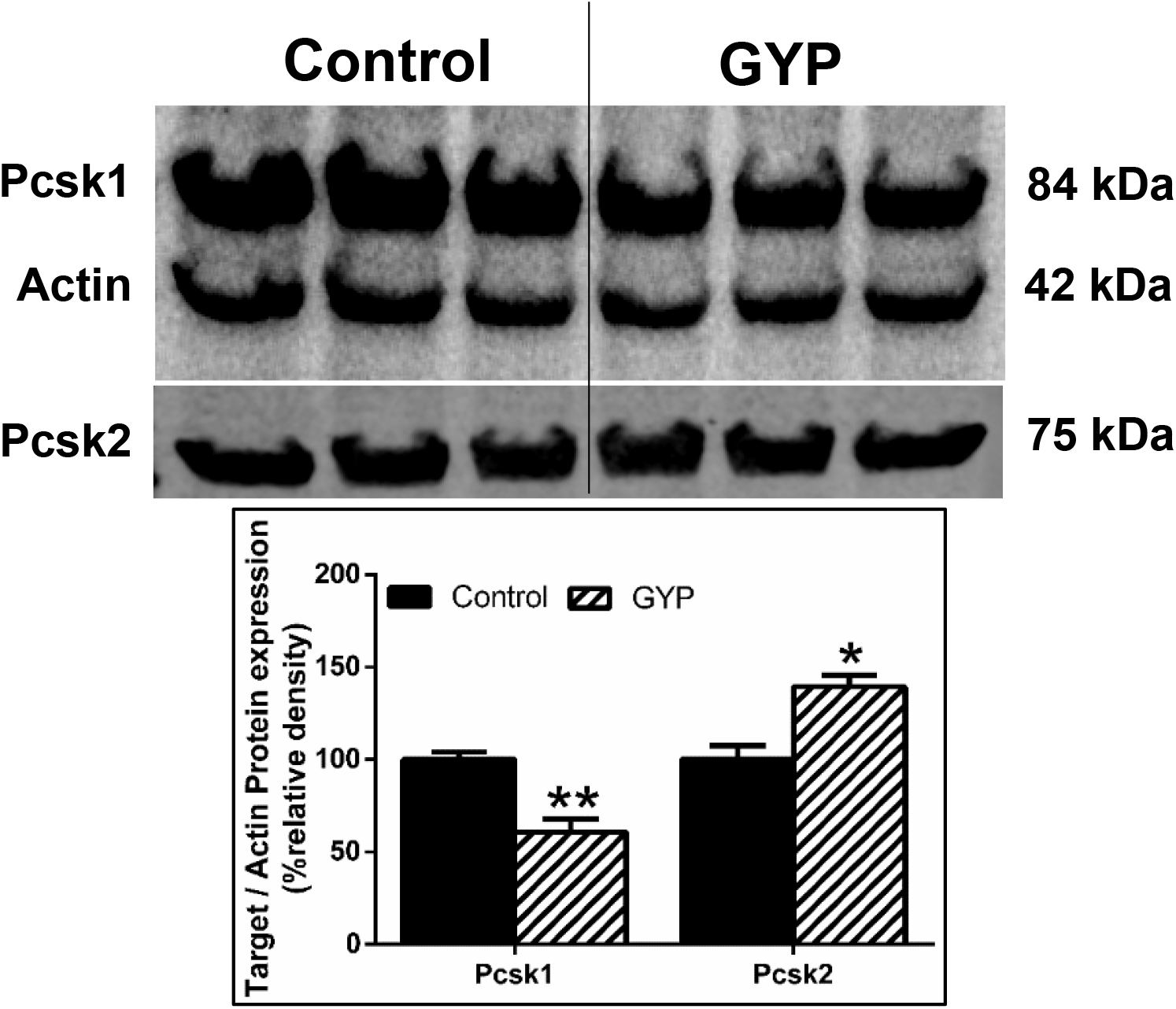
Effects of GYP treatment on Protein expression. Changes in Pcsk1 and Pcsk2 protein expression over 24 h treatment with GYP normalised to Actin expression. Plotted as %change to control. Values represent mean ±S.E.M. from three different experiments. Student’s t test was used for statistical analysis. *, P<0.05; **, P<0.01.

**Figure 6:**
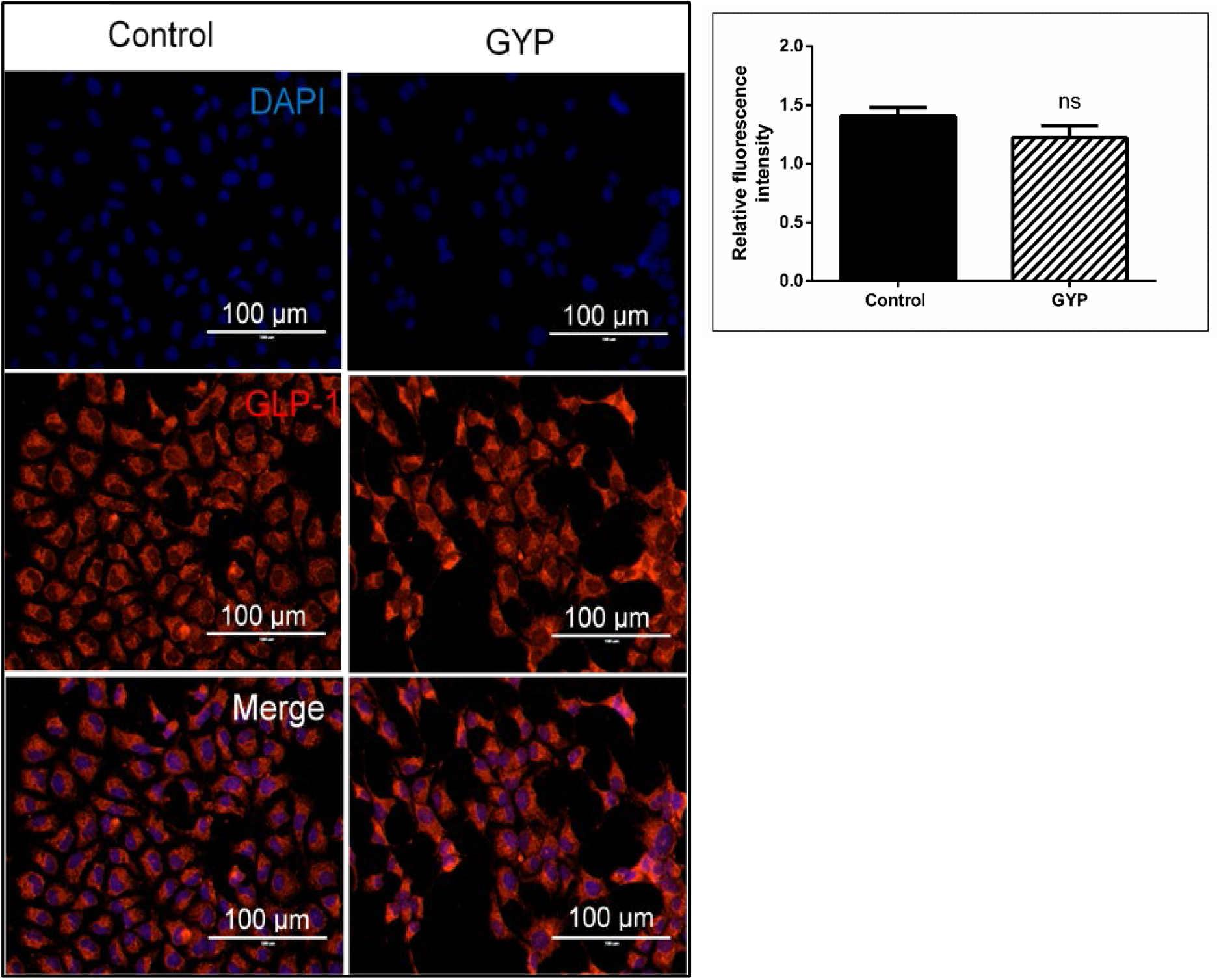
Effects of GYP treatment on GLUTag cellular GLP-1 content. GLUTag cells treated with 100μg/ml for 24h followed by staining with GLP-1 and nuclear stain DAPI. Fluorescence from control and GYP treated cells was measured (100 cells per coverslip from 4 separate experiments) and data was plotted as mean pixel intensity per 1000 pixel ^2^. Negative control was used to confirm specific GLP-1 staining and involved omission of primary anti-GLP-1 antibody only – samples without primary anti-GLP-1 stained positive for DAPI only.

## 4 Discussion

GLP-1 secreting L-cells are mostly concentrated along the distal part of the ileum and terminal colon. They are open-type intestinal epithelial cells that have contact with luminal nutrients via their apical surface while their basolateral surface is connected to underlying neural and vascular tissue (Baggio and Drucker, 2007). Both primary L-cells and the murine GLUTag L-cell model express sodium-glucose cotransporters 1 and 3 (Sglt1 and Sglt3) which co-transports two sodium ions with every molecule of glucose (Gribble et al., 2003; Reimann and Gribble, 2002). Previous studies in perfused rat small intestine and GLUTag cells confirmed that blocking Sglt1 completely abolished glucose-induced GLP-1 secretion (Gribble et al., 2003). A similarly impaired secretion was observed in perfused rat intestine when luminal Na^+^ was deprived, indicating Na^+^ coupled glucose uptake is necessary for GLP-1 secretion (Kuhre et al., 2015). The events following elevated intracellular Na^+^ via Sglt1 and GLP-1 secretion are not fully understood. It is believed that sodium/calcium exchangers (Ncx) play an important role in Na^+^ and Ca^2+^ homeostasis. The Ncx family exists in three subtypes Ncx1-3 encoded by Slc8a1-3 genes and they can operate either in the forward mode (Na^+^ entry and Ca^2+^ exit) or reverse mode (Na^+^ exit and Ca^2+^ entry) (Dong et al., 2005). Intestinal basal epithelial cells of small intestine, colon and rectum express Ncx1. Previous reports have shown that Vitamin D enhances Ncx1 gene expression and regulates transmembrane Ca^2+^ transport (Liao et al., 2019). In duodenal mucosal epithelium, bicarbonate secretion was regulated by Ncx1 reverse mode activation (Dong et al., 2005). A rat small intestinal epithelial cell line (IEC-18) expressing both Sglt1 and sodium/calcium exchanger family (Ncx1) has been shown to regulate intracellular sodium and calcium levels via activation of Ncx1 (Chen *et al.*, 2016). Perhaps a similar mechanism might help in maintaining L-cell membrane potential. In the present study, we have confirmed the gene expression of all three Ncx family subtypes Ncx1, Ncx2 and Ncx3 in GLUTag cells, although expression at the protein level was not investigated. However, the potential role of Ncx in sodium/calcium homeostasis and GLP-1 secretion in L-cells is yet to be established. In the present study, GYP stimulated calcium uptake acutely at both low and high glucose concentrations along with mRNA levels of both Sglt1 and Ncx1 indicating GYP might regulate intracellular Na^+^ and Ca^2+^ homeostasis in the GLUTag cells. Consistent with these calcium observations, GYP acutely enhanced GLP-1 secretion from GLUTag cells concentration-dependently, irrespective of glucose concentration. While these observations are limited to GLUTag cells in the current study, further investigation using primary L-cells isolated from human or animal models is warranted to establish the potential translation of these interesting findings. In pancreatic β-cells, gluco- and lipo-toxicity increase ROS generation and induces oxidative stress which can lead to cellular dysfunction and demise. Due to the very low expression of antioxidant enzymes which are necessary for neutralising ROS, oxidative stress causes β-cell apoptosis (Vasu *et al.*, 2015). Similar results were observed in GLUTag cells when exposed to elevated levels of glucose and fatty acid to induce cytotoxicity (Vasu *et al.*, 2015). Under conditions of oxidative stress, activation of the Nrf2/Keap1/ARE pathway is upregulated to enhance endogenous antioxidants which are necessary for free radical scavenging and cellular protection. Activation of superoxide dismutase genes (SOD1 and SOD2) by Nrf2 regulates intracellular superoxide levels by converting them to hydrogen peroxide, and Nrf2 mediated upregulation of catalase and glutathione peroxidase genes prevent hydrogen peroxide accumulation by converting it into water molecules (Pall and Levine, 2015). Ho-1 is another cytoprotective gene regulated by Nrf2 (Alam et al., 1999). It can be stimulated by several stimuli including cytokines, nitric oxide, heme, modified lipids and growth factors, and it acts to catalyse heme degradation to generate iron ion, biliverdin and carbon monoxide (CO), which are known for anti-apoptotic, anti-oxidant and anti-inflammatory properties (Loboda et al., 2016). In previous studies, Nrf2 activation promoted β-cell self-repair after high fat diet induced oxidative stress in rats (Abebe et al., 2017). The activators of Nrf2 have been shown to improve insulin sensitivity in mice and to protect against ROS and cytokine toxicity in human islet cells (Masuda et al., 2015; Yu et al., 2011). The role of Nrf2 pathway in β-cell protection is well characterised, although its function in intestinal L-cells is yet to be investigated.

In previous studies, GYP protected PC12 cells (a model of neuronal differentiation) from amyloid-beta induced apoptosis by activating Nrf2 pathway (Meng et al., 2014). Similar Nrf2 mediated protective effects against peroxide-induced toxicity have been reported in retinal epithelial cells (Alhasani et al., 2018). GYP treatment in a streptozotocin-induced diabetic rat model significantly upregulated Nrf2 expression and protected against oxidative stress (Gao et al., 2016). In agreement with these results, we have observed an increase in GLUTag cell viability following GYP treatment for 24h with upregulation of Nrf2 expression also noted in these cells along with upregulation of antioxidant Cat and Ho1 expression. Thus, GYP by upregulation of these antioxidant molecules in GLUTag cells may enable protection of the cells against increased cellular peroxide levels as may be observed under lipo and gluco-toxic conditions. The importance of this in *in vivo*, given the turnover of gut endocrine cells, requires further investigation in normal and obese/diabetes models.

The proglucagon gene is expressed in both intestinal L-cells and pancreatic α-cells, with post-translational processing of the propeptide by Pcsk1 generating GLP-1 in L-cells and Pcsk2 producing glucagon in α-cells (Drucker, 2006). It is known that the gut also produces extrapancreatic glucagon in humans (Lund et al., 2016). There is enough evidence that L-cells express and produce glucagon under specific conditions (Filippello et al., 2018; Kuhre et al., 2016). Chronic exposure to saturated free fatty acid palmitate caused increased glucagon and Pcsk2 expression, reduced Pcsk1 expression and GLP-1 secretion and might play a role in shifting L-cells towards glucagon production promoting glycaemic dysregulation (Filippello et al., 2018).

In this study, we found that acute exposure of GLUTag cells to GYP enhanced GLP-1 secretion irrespective of glucose concentration. However, prolonged 24h exposure of GLUTag cells to GYP resulted in a reduction in proglucagon mRNA, Pcsk1 protein levels and enhanced Pcsk2 mRNA and protein level. Although the mechanism underlying this reduction is unclear, it is not uncommon for reduced proglucagon expression following perhaps prolonged stimulation of GLP-1 secretion. A study by Lindqvist et al., (2017) showed that stimulatory concentrations of ghrelin enhanced GLP-1 secretion from GLUTag cells acutely yet following 24h treatment reduced proglucagon expression levels. While we noted a reduction in proglucagon gene expression, cellular staining intensity for GLP-1 did not appear to be reduced perhaps due to reserve GLP-1 granule pools, but this requires further investigation and quantification of GLP-1 content by ELISA.

## Conclusion

GLUTag cells express all 3 Ncx family ion channels and GYP treatment enhances Sglt1 and Ncx1 mRNA levels. Acutely, GYP induced calcium uptake and enhanced GLP-1 secretion from GLUTag cells irrespective of glucose concentration, while longer exposure of GLUTag cells to GYP downregulates proglucagon mRNA levels and up-regulates Pcsk2, although Pcsk1 mRNA and GLP-1 content were unchanged. Nrf2, Ho1 and Cat mRNA levels were upregulated in GLUTag cells following GYP treatment and enhanced cell viability. Further studies are required to confirm these results in primary L-cells and *in vivo* but suggest that GYP may modulate and enhance GLP-1 secretion acutely and protect cells against metabolic stress observed in obesity and diabetes.

## Abbreviations

GLP-1: Glucagon-like peptide-1
GYP: Gypenosides
T2DM: Type 2 diabetes mellitus
Ncx: Sodium/Calcium exchange channel
Sglt1: Sodium/glucose cotransporter 1
Gcg: Glucagon
Gck: Glucokinase
Pc1: Prohormone convertase-1
Pc2: Prohormone convertase-2
Nrf2: Nuclear factor erythroid 2-related factor 2
NFkb1: Nuclear factor Kappa b1
Cat: Catalase
Ho-1: Heme oxygenase-1
MTT: 3-(4,5-Dimethylthiazol-2-yl)-2,5-diphenyltetrazolium bromide

